# ILF3 contributes to the establishment of the antiviral type I interferon program

**DOI:** 10.1101/809608

**Authors:** Samir F. Watson, Nicolas Bellora, Sara Macias

**Affiliations:** Institute of Immunology and Infection Research, School of Biological Sciences, University of Edinburgh, King’s Buildings, Edinburgh, UK; IPATEC, CONICET, Bariloche, Argentina

**Keywords:** ILF3, NF90, NF110, Type I interferon, dsRNA, antiviral, translation

## Abstract

Upon detection of viral infections, cells activate the expression of type I interferons (IFNs) and pro-inflammatory cytokines to control viral dissemination. As part of their antiviral response, cells also trigger the translational shutoff response which prevents translation of viral mRNAs and cellular mRNAs in a non-selective manner. Intriguingly, mRNAs encoding for antiviral factors bypass this translational shutoff, suggesting the presence of additional regulatory mechanisms enabling expression of the self-defence genes. Here, we identified the dsRNA binding protein ILF3 as an essential host factor required for efficient translation of the central antiviral cytokine, *IFNB1*, and a subset of interferon-stimulated genes. By combining polysome profiling and next-generation sequencing, ILF3 was also found to be necessary to establish the dsRNA-induced transcriptional and translational programs. We propose a central role for the host factor ILF3 in enhancing expression of the antiviral defence mRNAs in cellular conditions where cap-dependent translation is compromised.

## Introduction

The presence of virus-derived double-stranded RNA (dsRNA) in the cytoplasm of infected cells is a hallmark of active viral replication. Mammals have developed several sensors, known as pathogen recognition receptors (PRRs), capable of recognising these virus-derived pathogen-associated molecular patterns (PAMPs). For instance, RIG-I-like receptors (RLRs), with MDA5 and RIG-I as central members of the family, sense dsRNA and signal through the mitochondrial associated antiviral factor, MAVS to induce the expression of type I interferons (IFNs) and proinflammatory cytokines (reviewed in (1)). Secreted type I IFNs bind the cell surface receptor IFNAR on infected and neighbouring cells to activate a second transcriptional response of approximately 500 interferon-stimulated genes (ISGs), which are responsible for establishing an antiviral state and preventing viral replication and dissemination (2)(3).

Virus-derived dsRNA also activates the cytoplasmic kinase PKR, which phosphorylates the initiation factor of translation eIF2α, resulting in a non-selective translational arrest of viral and cellular mRNAs also known as the host translational shutoff (4)(5)(6)(7). Regulation of eIF2α activity by phosphorylation provides a fast-acting mechanism to control protein expression in response to other stimuli, including amino acid starvation or endoplasmic reticulum stress (8)(9)(10). In addition to phosphorylating eIF2α and inducing the translational shutoff response, PKR can also phosphorylate the two major alternatively spliced isoforms encoded by the *ILF3* gene, known as NF90 and NF110 (11)(12)(13). NF90/110 are involved in regulating different steps of gene expression, including pre-mRNA splicing, miRNA biogenesis, and mRNA stability amongst others, and in controlling the life cycle of several viruses (reviewed in (14)(15)). Both ILF3 isoforms bind RNA through two tandem dsRNA-binding motifs and an RGG-rich domain (16)(17). In agreement with their association with polyribosomes, these factors can negatively regulate the translation of cellular mRNAs, and in particular, mRNAs containing AU-rich motifs (18)(19). In the context of viral infections, the current model suggests that NF90 and NF110 work in a complex with NF45 (20)(21)(22), and upon PKR-mediated phosphorylation, dissociate from NF45 and are retained on ribosomes to prevent translation of viral mRNAs (13). However, the translational targets of ILF3 during homeostasis or the antiviral response remain unknown. ILF3 isoforms have also been implicated in promoting the formation of stress granules during the antiviral response (23)(24), as well as being required for successful biogenesis of circular RNAs, a function that is impaired by activation of the antiviral response by the viral dsRNA mimic, poly (I:C) (25). However, the function of ILF3 during activation and establishment of the type I IFN response has not been characterized.

Besides the classical sensors of viral-derived dsRNA, other cellular dsRNA binding proteins are involved in limiting viral replication. Both dsRNA binding proteins TRBP and PACT regulate PKR activity and consequently the host-translational shutoff (26)(27). The OAS/RNase L system binds dsRNA and induces cleavage and degradation of RNA to limit viral replication, and participates in the translational shut off response by promoting turnover of the host mRNAs (28)(29)(30). In addition to their role as direct antiviral factors, the dsRNA binding proteins DICER, DGCR8 and DROSHA have shown to be essential to control the antiviral response (31)(32)(33)(34). Considering that both NF90 and NF110 can also bind dsRNA, and their previously reported role as direct antiviral factors by interfering with the function of viral-encoded proteins and viral RNAs, here we characterized the role of NF90/NF110 in regulating the activation of the IFN pathway by dsRNA stimulation and in the establishment of the host translational shutoff. By combining polysome profiling and high-throughput RNA sequencing analyses, we uncovered a role for NF90/NF110 in establishing the gene expression profile associated with the activation of the dsRNA-mediated type I IFN response, at the transcriptional and translational level. Specifically, the NF110 isoform was found to be essential for efficient translation of *IFNB1* mRNA, the central cytokine of the antiviral response, and a subset of ISGs in an environment where cap-dependent translation is compromised. In agreement, in the absence of NF90/NF110, cells displayed impaired antiviral activity, which correlated with attenuated production of ISGs. We propose a role for ILF3 in enhancing translation of *IFNB1* and ISGs during the host translational shutoff response, thereby providing effective levels of these antiviral proteins and ensuring a competent type I IFN response.

## Material and methods

### Cell lines, transfections, poly(I:C) and IFN-β stimulation

HeLa and A549 cell lines were maintained in DMEM supplemented with 10% (v/v) foetal calf serum (FCS) and 1% Penicillin/streptomycin at 37°C and 5% CO_2_. Transfections of poly(I:C) (2 μg/mL, HMW, tlrl-pic; Invivogen) were performed using Lipofectamine 2000 according to the manufacturer’s instructions. Four hours post transfection, cells were collected for downstream applications. For IFN-β stimulation, HeLa cells were treated with 100U of recombinant human IFN-β (Peprotech, 300-02BC) for 4 hrs and harvested. Knock-down experiments were performed by two consecutive rounds of transfection with siRNA pools against *ILF3* (L-012442-00-0005, Dharmacon) or *EIF2AK2* (L-003527-00-0005, Dharmacon) in HeLa or A549s. As a negative control, a non-targeting siRNA pool was used (D-001810-10-05, Dharmacon). Individual siRNAs against both major ILF3 isoforms were adapted from Guan et al., 2008 (35). For NF110 (Sense-CUACGAGAGCAAAUUCAA C[dT][dT], antisense–GUUGAAUUUGCUCUCGUAG[dT][dT]), and NF90 (sense-G[mC]CCACC[mU]UUG[2flC]UU[2flU]UUAU[dT][dT], antisense-AUAA[mA]AAGCAAAGGUGG[2flG]C) siRNAs were used. Briefly, cells were seeded at 50-60% confluency and transfected with 25 nM siRNAs using Dharmafect. After 24 hours, cells were split and transfected with a second round of siRNAs. Cells were collected for downstream processing 72 hours after the first transfection.

### RNA extraction and RT/qRT-PCR analyses

Total RNA was extracted using Trizol and reverse transcribed (RT) using MMLV (Promega) or Transcriptor Universal cDNA master (Roche) using random hexamers or oligo-dT and analysed by quantitative PCR (qPCR) in a Roche Lightcycler 480 II system. For oligonucleotides sequences see **Supplementary Table 1**. Analyses (2-ΔCt) were performed by normalization against *RN7SK* or 18S rRNA levels.

### Cell lysis and Western blotting

Cells were lysed in RIPA buffer (50 mM Tris-HCl, pH 7.5, 1% Triton X-100, 0.5% Na-deoxycholate, 0.1% SDS, 150 mM NaCl), supplemented with protease inhibitor cocktail (Roche), 5mM NaF and 0.2 mM Sodium orthovanadate. Protein lysates were mixed with reducing agent and LDS sample buffer (Novex, ThermoFisher) and denatured at 70 °C for 10 min and loaded in Novex Nupage 4-12% Bis-Tris gels. Gels were transferred onto nitrocellulose membrane using the iBlot2 system (ThermoFisher). Membranes were blocked with PBS-0.05% Tween and 5% milk for 1 hour at room temperature with agitation, before overnight incubation with primary antibodies. Antibodies against PKR (ab45427 Abcam), ILF3 (ab92355 Abcam), s6RP (2317S CST), eIF6 (3833S CST), ILF2 (ab154169 Abcam), α-tubulin (CP06 Merck), fibrillarin (ab5821 Abcam), phospho-eIF2α (Ser-51) (D9G8) (3398S CST), IFNAR1 (ab10739 Abcam), IFIT3 (ab76818 Abcam), OASL (ab191701), IRF1 (CST #8478), eIF3M (Bethyl, A305-029A), anti-rabbit HRP (CST) and anti-mouse HRP (Bio-Rad) were used. Proteins were visualised using ECL (Pierce) on a Bio-Rad ChemiDoc imaging system. Protein bands were quantified using ImageJ (v1.51p) software and normalized to α-tubulin or fibrillarin.

### ELISA, IFN bioactivity and Virus protection assay (TCID_50_)

A549 and HeLa cells were depleted of ILF3 and poly(I:C) stimulated as described above. Conditioned medium was harvested after 4 hr of poly(I:C) stimulation for HeLa cells and 6 hr in A549 to quantify IFN-β production by ELISA using the Quantikine human IFN-β ELISA kit (R&D systems, DIFNB0) according to the manufacturer’s protocol. Data were collected using a Varioskan Flash plate reader. For viral protection assays, A549 were transfected with either mock (non-targeting siRNA) or siRNAs against *ILF3* for 72 hours, prior to poly(I:C) transfection (2 μg/mL). Next, medium was collected, centrifuged for 3 minutes at 500xg, and filtered using a 0.22 μm filter. For ISG induction analyses, A549 cells were seeded and treated with a 1:2 dilution of conditioned medium for 4 hr and harvested for qRT-PCR analyses. For the 50% Tissue Culture Infective dose (TCID_50_) assays, A549 cells were seeded into 96 well plates and treated for 4 hr with the following conditioned medium dilutions: 1:2, 1:10, 1:50, 1:100 and 1:250. Next, cells were infected with eight serial dilutions of Echovirus 7, with at least 6 wells per dilution and incubated for at least 24 hours before counting infected wells. TCID_50_ values were calculated using the Spearman and Kärber algorithm, as in Witteveldt et al., 2019 (34). As a control, IFN-β signalling was blocked with 1.5 µg/mL of neutralizing antibodies against IFNAR1 for 2 hours prior to the conditioned media treatment, as previously described in Szabo et al., (36).

### Polysome Profiling

Gradient buffer solution (0.3 M NaCl, 150 mM MgCL_2_, 15 mM Tris-HCL pH 7.5, 2 mM DTT and 0.1 mg/mL cyclohexamide) containing different sucrose concentrations was layered in Beckman Coulter 14 mL polypropylene ultracentrifuge tubes. 0.5 mL of 60% sucrose followed by 1.6 mL of each, 50%, 42%, 34%, 26%, 18% and 10% sucrose solutions were poured followed by flash freezing in liquid nitrogen after every layer. Gradients were stored at −80°C until use and defrosted overnight at 4°C to allow them to equilibrate. For polysome gradient fractionation, one 10 cm plate of HeLa cells at 80 % confluency was used. Fresh medium was added to cells for 1 hr prior harvesting to ensure active protein synthesis. Next, 50 ng/mL cycloheximide (CHX, Sigma) was added to the medium and incubated for further 30 minutes at 37°C, before transferring to ice, where cells were washed twice with 2 mL ice-cold PBS containing 50 ng/mL CHX. To harvest, cells were scraped into 2 ml ice-cold PBS containing 50 ng/mL CHX and pelleted by centrifugation at 1400 rpm (300 x g) at 4 °C for 5 minutes. The supernatant was removed and 0.5 ml ice-cold lysis buffer (Gradient Buffer; 1% Triton X 100) was added, lysates were passed several times through a 25G needle and incubated on ice for 10 min. Lysates were then transferred to a 1.5 ml microfuge tube and centrifuged at maximum speed (20,000 x g) for 10 min at 4°C. Supernatant was next carefully layered onto a sucrose gradient, and centrifuged at 256,000 x g (38,000 rpm) for 1 hr 50 min at 4°C in a SW40 rotor in a Beckman L60 Ultracentrifuge. Ten 1 mL fractions were collected using a fraction collector (FoxyR1) monitoring optical density (Teledyne ISCO UA-6 detector with Optical unit 11). Constant flow rate was achieved using a pump syringe (Brandel) and Fluorinert FC770 (Flourinert). Optical absorbance of the solution was read at 254 nM and recorded directly using PeakTrak software V. 1.1. For protein extraction, 20% (v/v) final of cold TCA was added to each fraction and incubated on ice for 10 min prior centrifugation (20,000 x g for 5 min at 4 C). The pellet was washed twice in 1 ml acetone (−20°C) followed by centrifugation. After the last acetone wash, the pellet was air-dried and dissolved in 100 mM Tris-HCl pH 8.0 to be loaded in SDS-PAGE gels and western blot analyses. For RNA analyses, RNA was either extracted from each individual fraction or pooled into subpolysomal or polysomal pools. For this, three volumes of absolute ethanol were added to each fraction along with 0.1% SDS and precipitated overnight at −20°C. Samples were centrifuged at 5000 x g for 50 min at 4°C, and pellets resuspended in 50 μL water at 4°C with gentle agitation for an hour. Finally, Trizol LS was used to extract RNA, according to the manufacturer’s protocol. To disrupt polysomes by EDTA treatment, lysates were pre-treated with EDTA (50 mM) before sucrose fractionation.

For high-throughput sequencing analyses, RNA was extracted from pooled polysomal fractions (polysome profiling) or directly from cells (total RNA profiling). HeLa cells were either ILF3-depleted by ILF3-targeting siRNAs or transfected with non-targeting siRNAs as a control (siMock), followed by stimulation with poly(I:C). Experiments were performed in three biological replicates. For total RNA profiling, library preparation and sequencing for total RNA sequencing was carried out by BGI (Hong Kong). Briefly, total RNA was rRNA-depleted using the Ribo-zero rRNA depletion kit (Illumina) and strand-specific libraries were prepared using the Illumina Truseq protocol (Illumina) and sequenced on an Illumina HiSeq 4000 generating 100 bp paired end reads. For polysome profiling, library preparation and sequencing were carried out by Novogene (Hong Kong). Libraries were enriched for mRNAs using polyA + selection and non-strand-specific libraries prepared using the Illumina Trueseq protocol (Illumina) and sequenced on an Illumina HiSeq 4000 generating 150bp single end reads.

### Bioinformatic analyses

Total and polysome associated RNA-sequencing datasets were processed with the following RNA-seq pipeline; quantification of transcript and gene level abundances per sample were performed with Salmon v0.9.1 (parameters: --seqBias -g) (37) over the Gencode v24 reference human transcriptome (38). Estimated counts were retrieved as input for differential expression analyses. Only protein coding genes were considered in the case of polysomal samples. Statistical inference of regulated genes between each pair of conditions were calculated with the EgdeR v3.24.3 (39) R library using trimmed mean of M-value normalization (TMM) and the generalized linear model (GLM). Differentially regulated genes were examined with an FDR cut-off of 0.05 obtained with the Benjamini & Hochberg correction. MA-plots, heatmaps, boxplots were generated using gplots, pheatmap and the viridis R libraries. Overlapped MA-plot representations (**Figure 1B, 4A** and **4D**) were obtained using sliding bins along the complete range of the average expression logCPM (size = 1.5, step=0.5), quartiles and median were retrieved in bins with more than 25 genes. The average expression for each gene was calculated using the logCPMs obtained in each of the conditions compared.

**Figure 1.**
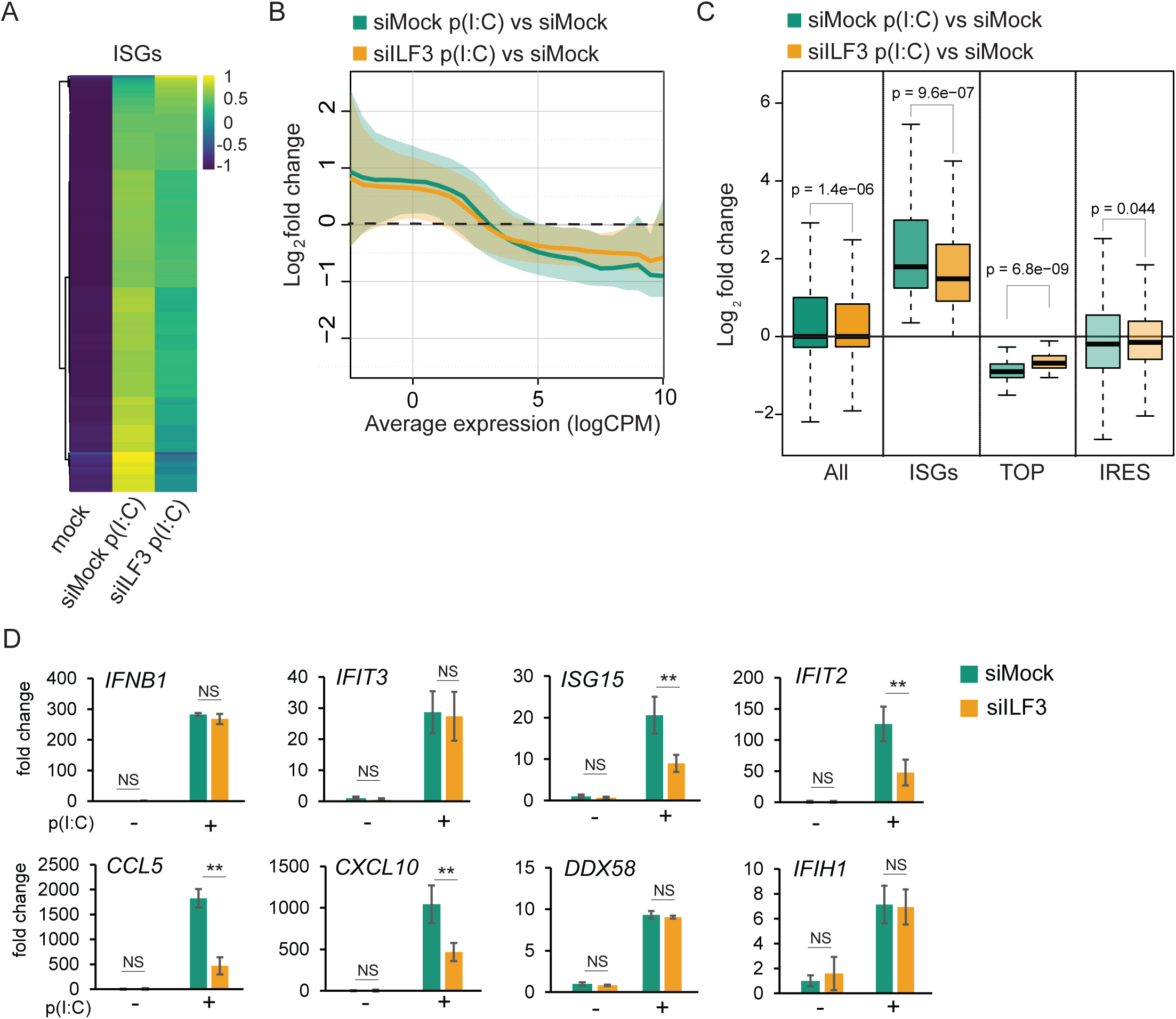
ILF3 is required to induce a robust type I IFN response. **(a)** Heatmap of significantly induced ISGs (from n=567 ISGs annotated in Interferome, n=374 were significantly induced after p(I:C) stimulation in HeLa cells), comparing expression values (CPM) normalised by gene **(b)** Log2 fold change of differentially expressed genes during dsRNA stimulation (siMock p(I:C) vs siMock, green) and during dsRNA stimulation in the absence of ILF3 (siILF3 p(I:C) vs siMock, yellow); solid line is the median value, and shaded area is the region between the first and third quartiles of the distribution. The average expression (x-axis) is calculated using the logCPMs for each gene obtained in the pairwise comparison **(c)** Box-plot analyses of differential gene expression during dsRNA stimulation for: all genes, significantly induced ISGs, TOP and IRES mRNAs in the presence (siMock p(I:C) vs siMock, green) or absence of ILF3 (siILF3 p(I:C) vs siMock, yellow), p-val by Mann-Whitney U test **(d)** qRT-PCR analyses of ISGs expression levels in the presence (siMock) or absence of ILF3 (siILF3), data show the average of at least 5 biological replicates ± std error, (**) pval<0.001 by two-way ANOVA followed by Tukey’s multiple comparison test.

For GC-content analyses, the longest isoform annotated for each protein coding gene was retrieved (Gencode v24, (38)). Genic regions were obtained according to the annotations and the following nucleotide contents calculated: GC-content (S/W), R/Y, A/T and G/C. To define the subset of GC or AT-richness, groups were chosen after splitting by the median of the total protein coding genes.

The list of ISGs was generated from the significantly upregulated genes in our total RNA-seq dataset and cross-referenced using the Interferome 2.0 database (40) with the following search conditions; Interferon type: Type I, Subtype: IFN-beta, Species: *Homo sapiens*. 5’TOP mRNAs were obtained from Yamashita et al., 2008 (41). IRES-containing genes were obtained from Weingarten-Gabbay et al., 2016 (42) and IRESite (www.IRESite.org) (43). For a complete list of ISGs, TOP and IREs genes see **Supplementary Excel File 1**.

Functional enrichment analysis of significantly differentially expressed genes were performed using the Kyoto Encyclopaedia of Genes and Genomes (KEGG, http://www.genome.jp/kegg/) and REACTOME (http://reactome.org) databases, implemented using the Intermine R library and the humanmine database (44)(45). KEGG and REACTOME enriched terms were selected using a false discovery rate threshold of 0.05 (Holm-Bonferroni correction).

## Results

### ILF3 depletion affects expression of the antiviral type I IFN program

During homeostasis, ILF3 has been involved in most steps of the gene expression pathway, from transcription to translation, including splicing, RNA stability and export (14). However, the role of ILF3 during cellular stress, such as the antiviral IFN response, is still unknown. To elucidate the function of ILF3 during the antiviral response, total gene expression analyses by RNA-sequencing were performed during ILF3 depletion in conditions of type I IFN activation (**Supplementary Figure 1A**). For this purpose, HeLa cells were depleted of ILF3 with siRNAs targeting both major isoforms, NF90 and NF110, and the type I IFN response was induced by transfection of the dsRNA analogue, poly(I:C). This analogue activates the expression of type I IFNs, but also induces translational shutoff by stimulation of PKR and RNAse L activity. Analyses of total RNA sequencing of poly(I:C)-stimulated HeLa cells confirmed the expression of the type I IFN *IFNB1*, but not *IFNA*, and induction of more than 300 classical interferon-stimulated genes (ISGs), as expected (**Figure 1A** and **Supplementary Excel File 2**). KEGG and Reactome analyses for upregulated genes were enriched for ‘Interferon alpha/beta signalling’ and ‘immune system’ categories, whereas metabolic genes were represented in the group of downregulated genes during dsRNA stimulation (**Supplementary Figure 1B**). The depletion of ILF3 significantly changed the levels of 12% of the poly(I:C)-induced genes and 10% of the poly(I:C)-downregulated genes, suggesting that most of the observed changes in gene expression upon activating the type I IFN response were conserved in the absence of ILF3 (**Supplementary Figure 2A** ‘siILF3 p(I:C) vs siMock p(I:C)’ and ‘siILF3 p(I:C) vs siILF3’), for depletion levels see **Supplementary Figure 2B**). During homeostasis, ILF3 depletion only caused a minor effect on the steady-state levels of transcripts, as less than 1% of the analysed genes displayed significant changes in their relative expression (**Supplementary Figure 2A** and **2C** for a complete list see **Supplementary Excel File 2**). A more detailed analysis of ILF3-dependent changes in gene expression revealed differential behaviour depending on the average expression of the genes (average expression is calculated using the logCPMs obtained for each gene in the two conditions compared). Lowly expressed genes were upregulated during dsRNA stimulation and were less increased in the absence of ILF3, whereas inhibited genes were highly expressed and less downregulated in the absence of ILF3 (**Figure 1B**). Specific analyses of annotated interferon-stimulated genes (ISGs) upregulated after poly(I:C) transfection revealed that ILF3 was necessary to stimulate their expression (**Figure 1A** and **1C**). Other subgroups of genes, such as IRES and TOP (5’terminal oligopyrimidine motif)-containing mRNAs were not negatively affected by ILF3 depletion, suggesting that the positive effect of ILF3 on expression was ISG-specific (**Figure 1C**). TOP mRNAs are enriched for genes encoding protein synthesis factors, such as ribosomal proteins, and their translation is regulated during stress growth conditions by the mTOR pathway (46). On the other hand, IRES-containing mRNAs bypass the need of 5’end cap structure to initiate translation and directly recruit the ribosome internally to the mRNA (47). These results were validated by qRT-PCR, confirming a reduction in the production of the ISGs, *IFTI2*, *ISG15*, *CCL5* and *CXCL10* in the absence of ILF3 (**Figure 1D**). The expression levels of *IFNB1* mRNA, the central player in initiating the type I IFN response and inducing the expression of ISGs, remained unchanged in the absence of ILF3 (**Figure 1D**). These analyses suggest that ILF3 is essential to induce a robust type I IFN gene expression program.

**Figure 2.**
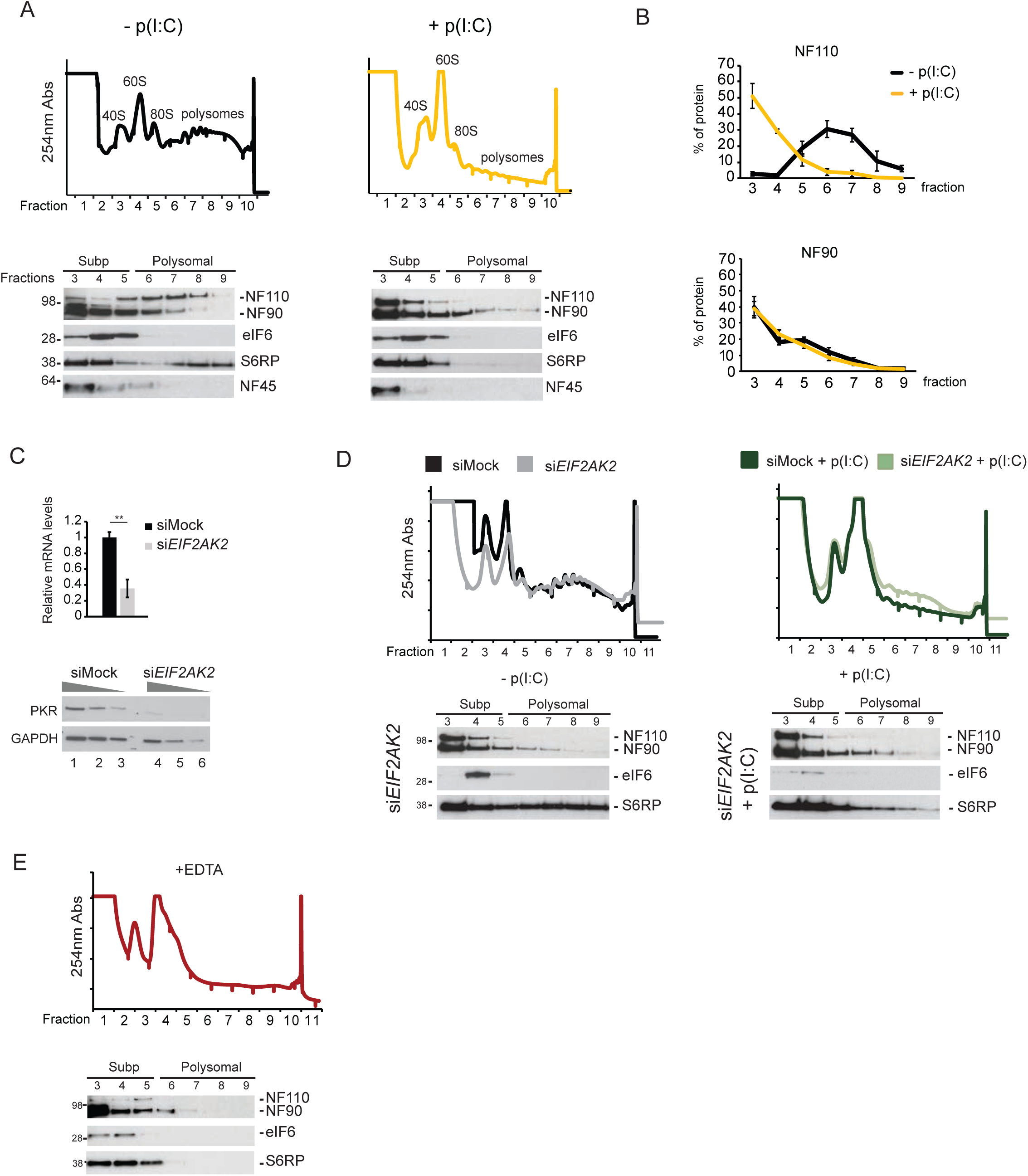
ILF3 isoforms associate with polyribosomes. **(a) (top)** Sucrose fractionation of cytoplasmic extracts from mock (top left) or poly(I:C) (top right) stimulated HeLa cells. UV absorbance (254 nm) is represented in the y-axis, for each of the fractions collected after centrifugation (x-axis) (**bottom**) Western blot analyses for co-sedimentation studies of NF110, NF90, and NF45 in the collected fractions. eIF6 and S6RP serve as markers for free 60S and mature ribosomes containing 40S, respectively (representative blots for each condition) **(b)** Average (n=5) distribution of the co-sedimentation of the NF110 (top) and NF90 (bottom) isoforms in polysomal fractionation during homeostasis (black) and the dsRNA-activated response (yellow**) (c)** qRT-PCR quantification of *EIF2AK2* mRNA after transient depletion in HeLa cells (top), data shown are the average (n=3) ± sem, (*) p-value < 0.05 by Student’s T-test. Western blot analyses of PKR protein levels upon transient depletion (bottom), GAPDH serves as a loading control (**d**) (**top**) Sucrose fractionation of cytoplasmic extracts from PKR (*EIF2AK2*) depleted HeLa cells during mock (left) or dsRNA-stimulated conditions (right) (**bottom**) Western blot analyses of NF90 and NF110 in fractions after gradient centrifugation of cytoplasmic extracts from PKR-depleted (si*EIF2AK2*) HeLa cells in the absence (left) and presence of dsRNA stimulation (right) (**e**) Sucrose fractionation of EDTA-treated cytoplasmic extracts.

### ILF3 associates with polysomes and regulates *IFNB1* mRNA translation

While the levels of some ISGs were reduced in the absence of ILF3, the mRNA levels of *IFNB1,* the major driver in inducing their expression, were unchanged. Considering the previously reported role of ILF3 on mRNA translation (18) (19), we next aimed to study if ILF3 could be regulating the translation of *IFNB1* mRNA, and as consequence differential ISG expression. To test this possibility, we assessed the rate of *IFNB1* translation by polysome profiling, in addition to measuring IFN-β protein levels by ELISA. First, co-sedimentation of the different ILF3 isoforms with polysomes was characterized by sucrose fractionation analyses of HeLa cytoplasmic extracts stimulated with or without poly(I:C). During homeostasis, both major alternatively spliced isoforms of *ILF3*, NF90 and NF110, were found to associate with polysomes (**Figure 2A**, fractions 6-9), as well as lighter fractions, containing the 40S and 60S ribosomal particles (**Figure 2A** fractions 3-4, respectively), and monosomes (80S) (**Figure 2A**, fraction 5, quantification in **Figure 2B**). After stimulation with poly(I:C), a major drop in the levels of actively translating polysomes was observed, as expected by the dsRNA-activated translational shutoff (**Figure 2A**, top right panel). The translational shutdown was further confirmed by assessing the co-sedimentation profiles of ribosomal markers. eIF6 serves as a marker for pre-ribosomal subunit 60S, which dissociates from the nascent 60S particle when the mature 80S ribosome is formed (48). The ribosomal protein S6 (S6RP) serves as a marker for the small 40S ribosomal subunit. In poly(I:C)-stimulated conditions, S6RP no longer co-sedimented in the heavier polysomal fractions, confirming successful activation of the translational shutoff response (**Figure 2A**, bottom right panel). Under these conditions, NF110 isoform disengaged from heavier polysomal fractions, accumulating in the lighter fractions (**Figure 2A**, quantification in **Figure 2B** top panel), whereas NF90 profile remained similar when compared to non-stimulated cells (**Figure 2A**, quantification in **Figure 2B** bottom panel).

Both NF90 and NF110 are known substrates of the kinase PKR, one of the essential factors driving the translational shutoff during the dsRNA-mediated antiviral response (13). We therefore measured ILF3 isoforms association with polysomes in the absence of PKR (**Figure 2C**). Interestingly, the NF110 isoform shifted to lighter molecular weight fractions in the absence of PKR during homeostasis (**Figure 2D** left panel, compared to **Figure 2A** left panel), suggesting that association of this isoform with polysomes depends on PKR. Stimulation of PKR-depleted cells with poly(I:C) did not result in further changes in NF90 and NF110 sedimentation, when compared to non-stimulated PKR-depleted cells (**Figure 2D**), even though a less pronounced translational shutoff was observed, as evidenced by residual S6RP co-fractionation with higher molecular weight fractions (compare **Figure 2D** right panel with **Figure 2A** right panel). To confirm that ILF3 isoforms were indeed associated with polysomes, cytoplasmic extracts pre-treated with EDTA were fractionated to monitor NF90/NF110 sedimentation. EDTA treatment, which forces dissociation of ribosomal subunits, resulted in both NF90 and NF110 disengaging from the polysomal fractions (**Figure 2E**, from fraction 6 onwards). All these data support that NF90 and NF110 directly associate with actively translating ribosomes, and NF110 association with polysomes is dependent on PKR.

Considering these results, we next aimed to determine if ILF3 could regulate *IFNB1* translation. We performed polysomal fractionation and assessed *IFNB1* mRNA association with polysomes as an indirect measurement of its translation, in the presence or absence of ILF3. We again confirmed that ILF3 depletion did not significantly affect the levels of *IFNB1* mRNA induction (**Figure 3A**). In addition, we confirmed that depletion of NF90/NF110 did not change the levels of eIF2α phosphorylation, but resulted in destabilization of NF45 (**Figure 3B**), as previously reported (35). We next studied the sedimentation of *IFNB1* mRNA with polysomes in the presence or absence of ILF3 by semi-quantitative RT-PCR on each of the collected fractions, as in **Figure 3C**. Successful poly(I:C)-activated translational shutoff was confirmed by the shift of *FOS* mRNA from heavier to lighter polysomal fractions (**Figure 3D**, compare top left and right panels). Interestingly, the depletion of ILF3 affected the distribution and association of *IFNB1* mRNA with polysomes, suggesting that ILF3 regulates translation of this mRNA (**Figure 3D**, bottom gels). In addition, *IFNB1* mRNA in pooled subpolysomal and polysomal fractions was quantified by qRT-PCR and confirmed that ILF3 depletion significantly decreased the co-sedimentation of *IFNB1* mRNA with polysomes, suggesting that ILF3 may indeed regulate IFN-β protein production (**Figure 3E**). This was confirmed by quantifying IFN-β protein levels by ELISA in the supernatants of both HeLa and A549 cells after stimulation with poly(I:C). ILF3 depleted cells showed a significant decrease in IFN-β protein levels (**Figure 3F**).

**Figure 3.**
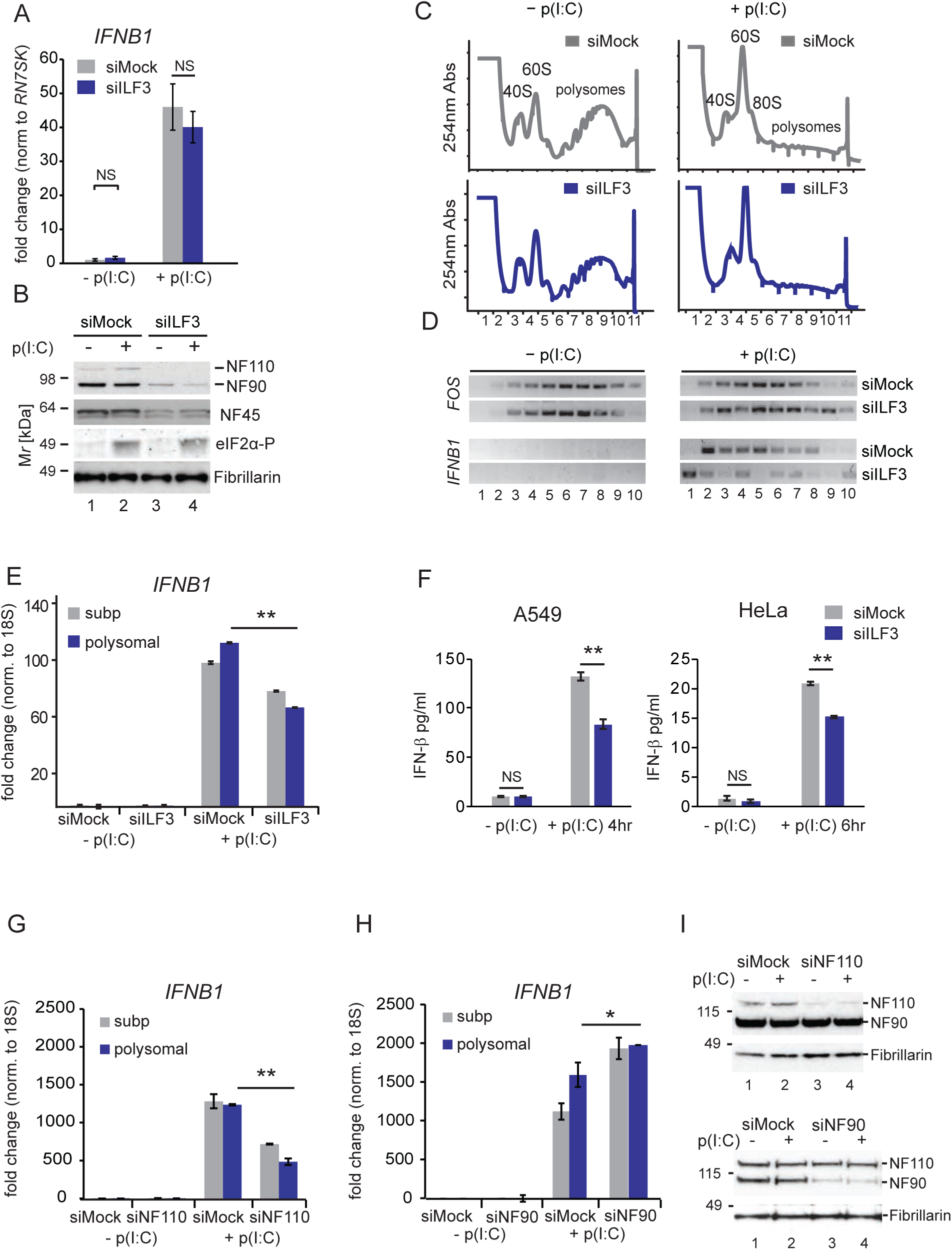
NF110 is necessary for *IFNB1* mRNA translation. (**a**) Quantification of *IFNB1* mRNA levels 4 hours after poly (I:C) transfection in mock (siMock, grey) or ILF3-depleted cells (siILF3, blue) by qRT-PCR. Data show the average (n=12) ± sem relative to mock and normalized to *RN7SK*. p-val (N.S., not significant by two-way ANOVA followed by Tukey’s multiple comparison test (**b**) Representative western blot analyses for NF90 and NF110 depletion after siRNA transfection, and NF45 and phosphorylated eIF2α levels. Fibrillarin serves as a loading control (**c**) (**top**) Sucrose fractionation of cytoplasmic extracts from mock siRNA transfected HeLa cells (grey) stimulated or not with poly(I:C) (top right vs top left). (**bottom**) Polysomal fractionation of siILF3 depleted HeLa cell extracts (blue), stimulated or not with poly(I:C) (right vs left). UV absorbance (254 nm) is represented in the y-axis, for each of the fractions collected after centrifugation (x-axis) (**d**) RT-PCR detection of *FOS* (top) and *IFNB1* (bottom) mRNA co-sedimentation in each of the fractions collected in (**c**) (**e**) qRT-PCR analyses of *IFNB1* mRNA relative abundance in subpolysomal and polysomal pooled fractions. Data show the average (n=3) ± sem normalised to 18S rRNA and relative to subpolysomal levels in mock, (**) pval<0.001 by two-way ANOVA followed by Tukey’s multiple comparison test (**f**) IFN-β quantification by ELISA from poly(I:C) stimulated A549 cells (left) and HeLa (right) upon ILF3 depletion, compared to mock-depleted cells (siMock). Data show the average (n=4 in A549, n=3 in HeLa ± sem, (**) pval<0.001 by two-way ANOVA followed by Tukey’s multiple comparison test (**g, h**) qRT-PCR quantification of *IFNB1* mRNA relative abundance in subpolysomal and polysomal pooled fractions from NF110 (**g**), or NF90 depleted HeLa cells (**h**), data show the average,(n=3 in (**f**), n=2 in (**g**)) ± sem, normalised to 18S rRNA and relative to subpolysomal levels in mock, (*) pval<0.05, (**) pval<0.001 by two-way ANOVA followed by Tukey’s multiple comparison test, N.S. not significant (**i**) Representative western blot analyses for NF110 (top) and NF90 (bottom) isoform depletions for experiments in (**g)** and (**h)**, respectively. Fibrillarin serves as a loading control.

As *ILF3* encodes the two major isoforms NF90 and NF110 and only the NF110 association with polysomes was affected during poly(I:C) stimulation, we next assessed if regulation of *IFNB1* mRNA association with polysomes is isoform specific. For this purpose, we designed specific siRNAs targeting NF90 or NF110. Extracts depleted for these factors were fractionated, and subpolysomal and polysomal fractions were pooled to quantify the *IFNB1* mRNA in these two populations. Interestingly, the depletion of NF110, but not NF90, caused a decrease in the association of *IFNB1* mRNA with the polysomal fractions (**Figure 3G-3I**). All these results suggest that translation of *IFNB1* mRNA is enhanced by the NF110 isoform of ILF3 after dsRNA stimulation.

### ILF3 is necessary for the host translational shutoff and translation of ISGs

Considering the previously reported role of ILF3 in regulating translation during homeostasis and its role in enhancing *IFNB1* mRNA translation, we next wanted to characterize the impact of ILF3 function in global translation during the type I IFN response. For this purpose, mRNA was extracted in triplicates from pooled polysomal fractions from poly(I:C) stimulated and unstimulated HeLa cells in the presence or absence of ILF3 and the transcriptome was sequenced (**Supplementary Figure 3A**). After dsRNA stimulation, we observed two populations of protein-coding mRNAs that were differentially affected by ILF3. First, a population of low abundance transcripts that became more associated with polysomes during dsRNA stimulation, and second, a highly expressed population that became less associated (**Figure 4A** and **Supplementary Figure 3B**, for complete list see **Supplementary Excel File 3**). Interestingly, those mRNAs that were recruited to polysomes during activation of the type I IFN response, which were mainly cytokines and ISGs, were less engaged in the absence of ILF3 (**Figure 4A**, for KEGG analyses **Supplementary Figure 4A**). Conversely, highly associated mRNAs, which contain housekeeping genes and ribosomal proteins, no longer dissociated from polysomes during poly(I:C) stimulation upon ILF3 depletion (**Figure 4A**, for KEGG analyses **Supplementary Figure 4B**). To explore these observations further, we analysed subgroups of protein-coding genes separately and confirmed that ISGs were significantly less enriched in polysomes in the absence of ILF3, whereas TOP mRNAs became more enriched, and IRES mRNAs remained similarly enriched in polysomes upon ILF3 depletion (**Figure 4B**). There are several features in mRNAs that dictate their translatability, for instance, the GC-content of an mRNA positively correlates with efficient polysomal association (49). In agreement with this, we observed that GC-rich genes were also more associated with polysomes upon dsRNA stimulation, whereas AT-rich genes were disengaged, and this effect was attenuated upon ILF3 depletion (**Figure 4C**). A more refined analysis, considering the average expression of these genes, revealed that ILF3 depletion reverted some of these observations. Highly expressed AT-rich genes were no longer disengaged from polysomes, and low and medium-expressed GC-rich genes were less efficiently associated with polysomes (**Figure 4D**). Importantly, ISGs have a heterogeneous GC-content distribution, suggesting that the differential association of these genes with polysomes cannot be directly attributed to the GC-content effect revealed by our analyses.

**Figure 4.**
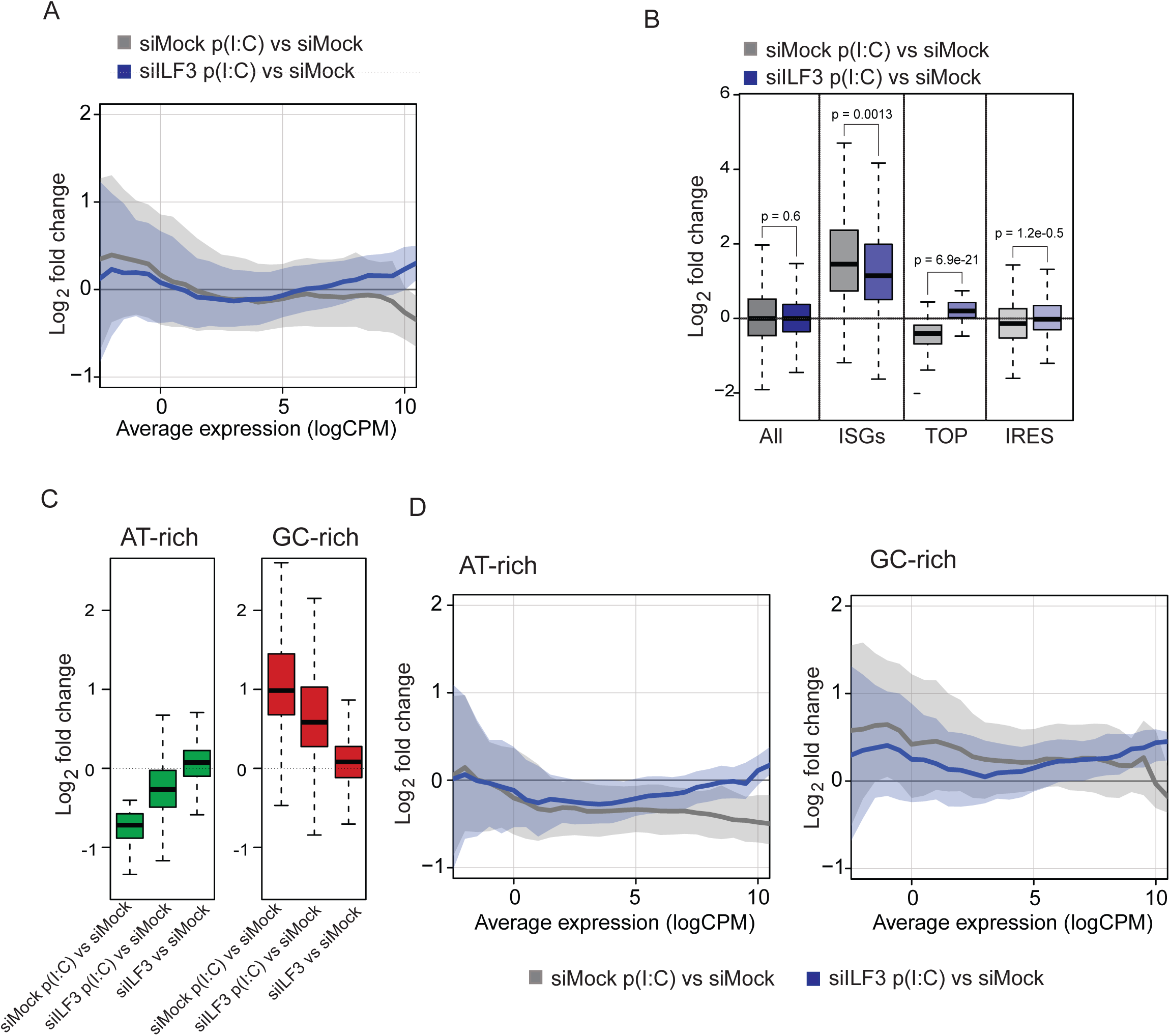
ILF3 controls dsRNA-mediated translational response. **(a)** Log2 fold change of differential enrichment of genes in polysomal fractions during dsRNA stimulation (siMock p(I:C) vs siMock, grey) and in the absence of ILF3 (siILF3 p(I:C) vs siMock, blue), x-axis represents the average expression for each of the genes in the two compared conditions (**b**) Box-plot analyses of differential enrichment in polysomal fractions for: all genes, induced ISGs, TOP and IRES mRNAs during the type I IFN response (siMock p(I:C) vs siMock, grey) and upon ILF3 depletion (siILF3 p(I:C) vs siMock, blue), Mann-Whitney U test p-value (**c**) Box-plot analyses of differential enrichment in polysomal fractions for AT-rich (left) and GC-rich (right) genes upon the antiviral response (siMock p(IC) vs siMock) and after ILF3 depletion during the antiviral response (siILF3 p(IC) vs siMock) and homeostasis (siILF3 vs siMock) (**d**) Log2 fold change of differential enrichment of AT- and GC-rich genes in polysomal fractions during dsRNA stimulation (siMock p(I:C) vs siMock, grey) and in the absence of ILF3 (siILF3 p(I:C) vs siMock, blue), x-axis represents the average expression for each of the genes in the two compared conditions.

Our data suggest that the ISGs and TOP mRNAs subgroups display opposite behaviours in polysomal association upon ILF3 depletion. ISGs association with polysomes was affected by the absence of ILF3 (**Figure 5A**), whereas TOP mRNAs, which are disengaged from polysomes upon activating the host translational shutoff, reverted to almost homeostatic levels in the absence of ILF3 (**Figure 5B**). qRT-PCR analyses of pooled subpolysomal and polysomal fractions confirmed that ILF3 depletion led to a significant reduction in polysome association for the ISGs *IFIT3, CCL5, IFIT2, ISG15, CXCL10* and *DDX58,* supporting a positive role of ILF3 in ISGs translation (**Figure 5C**). At the protein level, we confirmed that in the absence of ILF3, ISGs such as IFIT3, OASL and IRF1 were less induced (**Figure 5D**). The opposite effect was confirmed for the TOP mRNA, *EIF3M*. Polysomal association of *EIF3M* was lost during dsRNA stimulation, and this effect was reversed by depletion of ILF3, as a control *ACTB* association was quantified (**Supplementary Figure 5A** and **5B**). However, no changes in eIF3M protein levels were observed, which may be due to the its high stability, and the brief time of stimulation with poly (I:C) (**Supplementary Figure 5A**).

**Figure 5.**
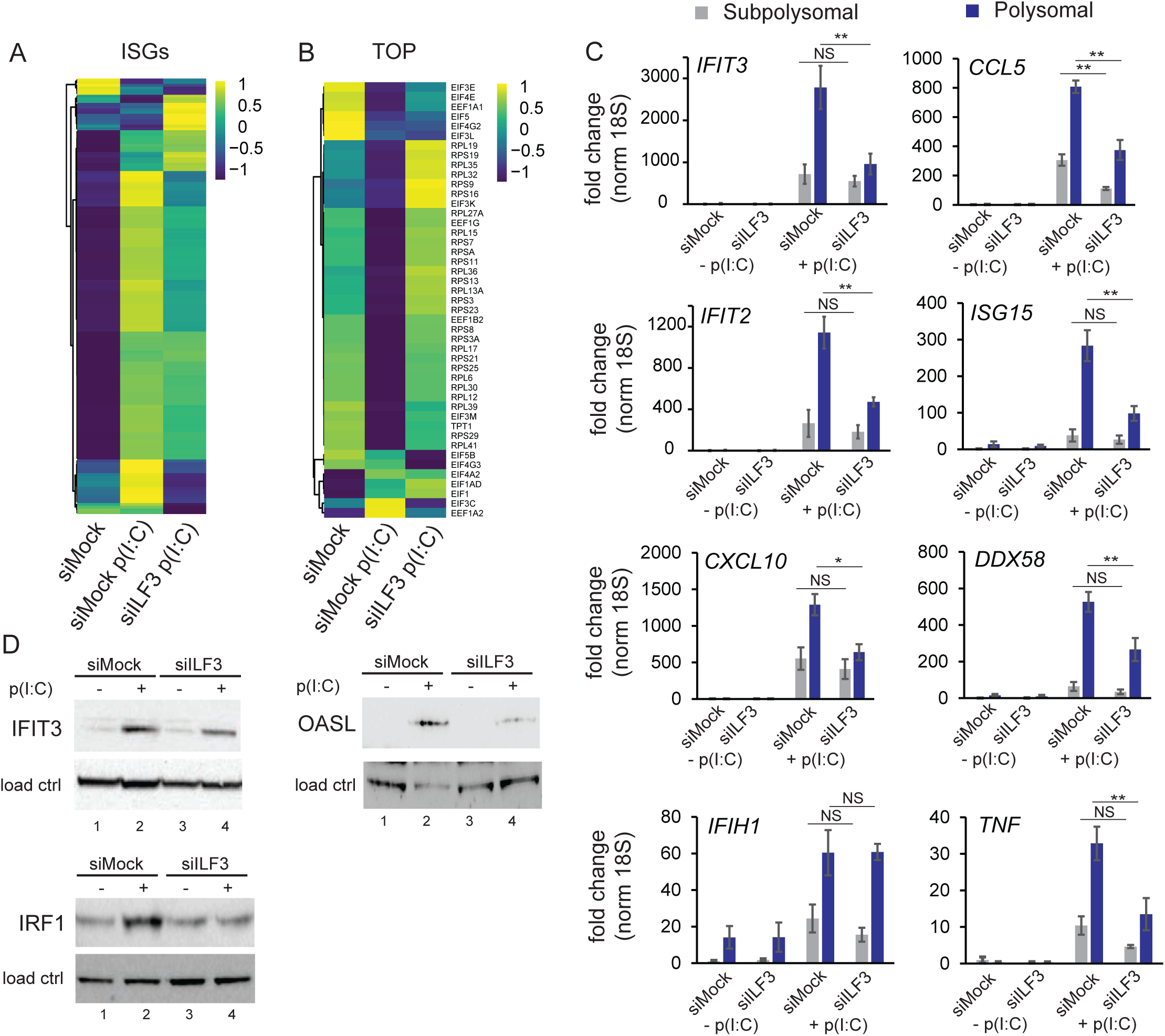
ILF3 is necessary for translation of ISGs. (**a**) Heatmap of ISGs enrichment in polysomal fractions (n=374) during homeostasis (mock), versus activated type I IFN response (siMock p(I:C)) and activated IFN response in the absence of ILF3 (siILF3 p(I:C)) (**b**) Heatmap of TOP mRNAs enrichment in polysomal fractions (n=45) in the same conditions as in (**a**) (**c**) qRT-PCR quantification of ISG enrichment in subpolysomal (grey) and polysomal (blue) pooled fractions in mock (-p(I:C)) or stimulated (+p(I:C)) HeLa cells, in the presence or absence of ILF3. Data show the average (n=3) +/− s.e.m, normalised to 18S rRNA and relative to subpolysomal level in mock (*) pval<0.05, (**) pval<0.001 by two-way ANOVA followed by Tukey’s multiple comparison test, N.S, non-significant (**d**) IFIT3, OASL and IRF1 western blot analyses upon dsRNA stimulation (lane 2), and in the absence of ILF3 (lane 4), tubulin and fibrillarin serve as loading controls.

To further elucidate the function of ILF3 on ISGs mRNA translation, we assessed if differential ISG polysomal association could be directly attributed to ILF3 presence or be a secondary consequence of lowered IFN-β production and ISG expression. To this end, polysome profiles of cells depleted or not of ILF3 and stimulated with recominant IFN-β were compared. A proportion of the tested ISGs (*IFIT3*, *IFIT2* and *ISG15*) followed the same behaviour as seen after poly(I:C) stimulation, suggesting that the differential polysomal association of these ISGs were due to direct ILF3 action (**Supplementary Figure 6**).

**Figure 6.**
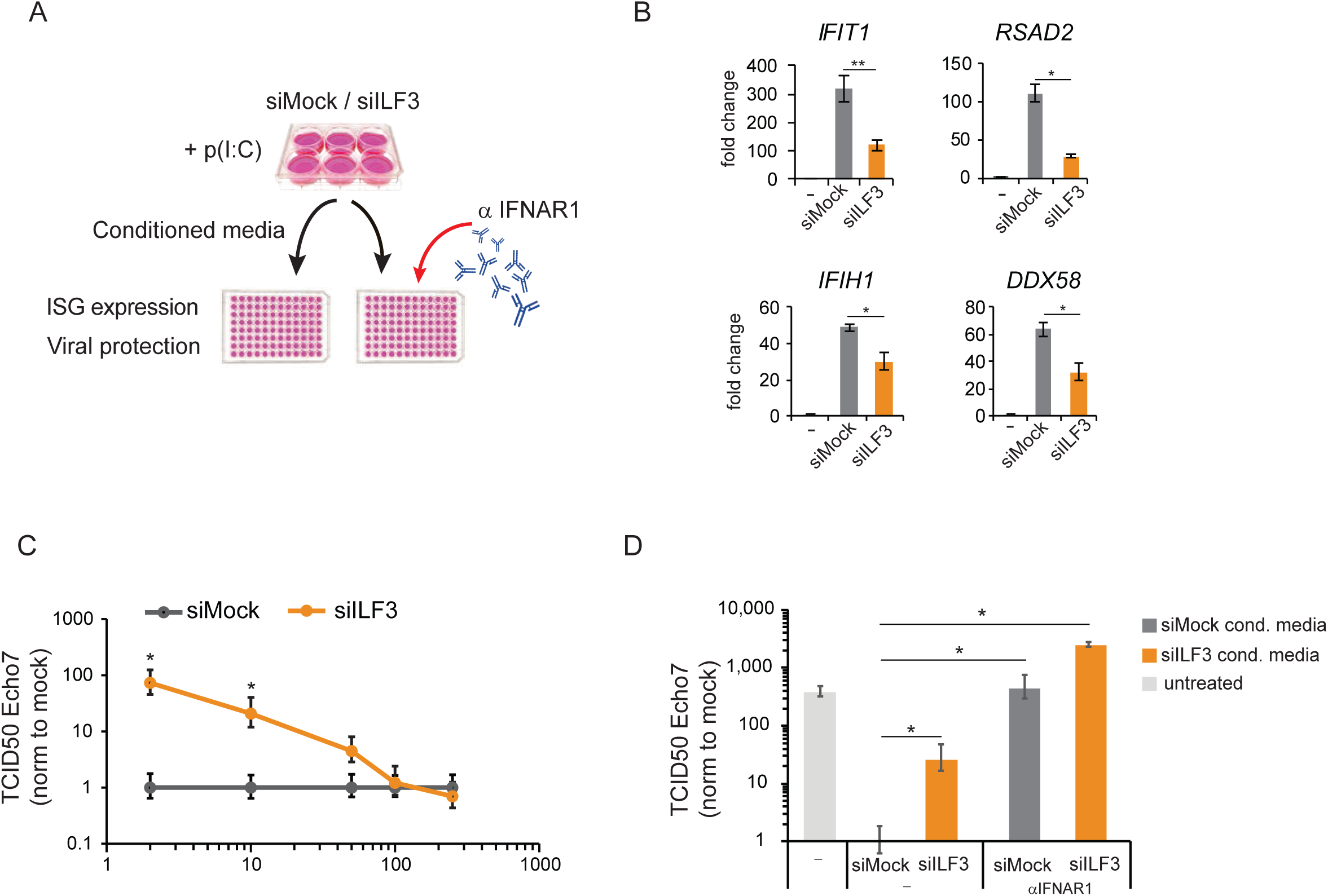
ILF3 is necessary to establish antiviral protection. **(a)** Schematic representation of the experiments performed in (**b**), (**c**) and (**d**). A549 cells were siMock or ILF3-depleted followed by stimulation with poly(I:C). Medium was collected, and specific or serial dilutions were added into fresh A549 cells to test ISGs induction levels or conferred protection to viral infections **(b)** qRT-PCR quantification of ISG expression levels after incubation with siMock (grey) or ILF3-depleted conditioned medium (yellow). Data show the average (n=4) ± sem, (*) pval<0.05, (**) pval<0.001 by one-way ANOVA followed by Tukey’s multiple comparison test **(c)** TCID_50_ assay using Echovirus 7 on A549 cells pre-incubated with serial dilutions of conditioned medium (y-axis) from ILF3-depleted (yellow) or siMock-depleted A549 cells. Data show the average (n=3) ± sd normalized to each siMock dilution, (*) p-value < 0.05 by Student’s T-test **(d)** TCID_50_ assay using Echovirus 7 on A549 cell pre-incubated with a single dilution (20x) of conditioned medium produced in siMock (dark grey) or ILF3-depleted cells (yellow) and, as a control, non-conditioned medium was added (light grey). Prior to infection, A549 were pre-incubated with anti-IFNAR1 receptor antibody. Data show the average (n=3) ± sd, (*) p-value < 0.05 by Student’s T-test.

All these data suggest that ILF3 has a dual function on mRNA translation during the type I IFN response by enabling translation of ISGs mRNAs and impairing TOP mRNA polysomal association. In addition, ILF3 was found to repress association of AT-rich genes with polysomes. All these together support the hypothesis that ILF3 has a major role in regulating the association with polysomes of important subtypes of genes during the host translational shutoff.

### ILF3 is necessary for type I IFN induction in response to dsRNA

As we have shown that ILF3 has an essential role in the expression of ISGs, we next wanted to evaluate the functional relevance of this regulation. For this purpose, mock or ILF3-depleted A549 cells were stimulated with poly(I:C) and the supernatant, or conditioned medium, was collected and added to naive cells to measure ISG induction and confirm differential type I IFN production and secondly, measure differential protection against viral infections (**Figure 6A**). Addition of conditioned medium from poly(I:C) stimulated cells induced the expression of ISGs *IFIT1*, *RSAD2*, *IFIH1* and *DDX58*, and as expected, medium from ILF3-depleted cells resulted in significantly lower expression levels of the same ISGs (**Figure 6B**), confirming that the production of type I IFN, which is essential to drive the expression of these genes, is diminished in the absence of ILF3.

As a functional approach, we tested if conditioned medium generated in the absence of ILF3 conferred decreased resistance to viral infections. Serial dilutions of conditioned medium were added to fresh A549 cells to test their sensitivity to Echovirus 7 infection by TCID_50_. Conditioned medium generated in ILF3-depleted cells was able to confer less resistance to Echovirus 7 infections, with an almost 100-fold reduction in protection observed at the lowest dilution (**Figure 6C**). To verify if the differential expression of IFN-β was in part responsible for the observed difference in susceptibility, one of the subunits of the type I IFN receptor, IFNAR1, was neutralized by pre-incubating cells with an anti-IFNAR1 antibody before addition of conditioned medium. Mock versus ILF3-depleted conditioned medium resulted again in significant differences in conferring viral protection to Echovirus 7 and blocking of IFNAR receptor impaired the antiviral protection effect, rendering cells susceptible to Echovirus 7 reaching similar levels to the untreated control (**Figure 6D**). These results support the hypothesis that differences in viral protection observed between mock and ILF3-depleted cells are caused by differential production of ISGs, highlighting again the central role for ILF3 in facilitating the establishment of a robust cellular type I IFN response upon dsRNA stimulation.

## Discussion

The activation of the dsRNA-mediated antiviral response involves drastic changes in the gene expression program of cells which needs to be tightly regulated from transcription to translation. In this work, we propose that the RNA-binding protein ILF3, which is involved in many steps of RNA processing during homeostasis, is also essential for establishing a robust type I IFN program upon dsRNA stimulation. We found this factor acts at the level of induction of ISGs as well as enhancing their translation. These results agree with the previously reported antiviral role of the ILF3 isoforms in the context of HIV and (+) ssRNA viral infections, although these effects were proposed to be mediated by direct binding of ILF3 isoforms to the viral genome or viral proteins (50)(51)(52)(53). To assess whether ILF3-mediated regulation of the gene expression pathway was mediated by direct association of ILF3 with the different transcripts, we compared our results to publicly available CLIP datasets (54). Unfortunately, ILF3 binding is too ubiquitous to obtain clear correlations between binding and changes in gene expression upon ILF3 depletion. Whereas our analyses revealed that ILF3 binds to thousands of different genes, it primarily overlapped with Alu-derived sequences (data not shown), confirming similar recent observations (55)(25).

ILF3 had been previously suggested to have an inhibitory role for the translation of mRNAs harbouring AT-rich motifs during homeostasis (18)(19). Interestingly, our results suggest a similar role for ILF3 during the type I IFN response. AT-rich mRNAs were dissociated from polysomes during the host translational shutoff, but this effect was reverted in the absence of ILF3, confirming a role of ILF3 in inhibiting translation of AT-rich mRNAs during the host translational shutoff. In addition, ILF3 was found to be essential for efficient polysomal dissociation of TOP mRNAs upon dsRNA stimulation. This effect did not seem to be a consequence of differential activation of the host translational shutoff in the absence of ILF3, since similar levels of phosphorylated eIF2α were detected. However, a decrease in total mRNA levels for TOP genes was also observed upon dsRNA stimulation (**Figure 1C**), suggesting that the differential polysome loading could be the combined result of regulating the available pool of TOP mRNAs as well as their association with polysomes. In contrast to these, ILF3 was shown to be required for efficient translation of *IFNB1* and a subgroup of ISGs, suggesting an additional role for ILF3 as a positive factor for translating essential self-defence genes during the inhibition of cap-dependent translation. Regarding ISGs, such as *IFIT3* and *DDX58*, ILF3 only affected their association with polysomes, resulting in decreased protein production. However, in a considerable proportion of the ISGs analysed, ILF3 depletion resulted in both changes in their RNA steady state levels and their polysomal association, suggesting that both effects could provide a certain level of redundancy to guarantee optimal levels of these antiviral factors. An additional layer of complexity is provided by the fact that ILF3 directly regulates the production of the main inducer of ISGs, the type I IFN, IFN-β. To disentangle the specific contribution of ILF3 in controlling ISGs translation, polysome profiling of these mRNAs was studied upon exogenous IFN-β stimulation in the absence of ILF3. These analyses revealed that the role of ILF3 in controlling ISG levels and polysome loading was largely conserved between dsRNA and exogenous IFN-β stimulation, suggesting that direct ILF3 action and not the differential expression of IFN-β is responsible for this regulation. Considering these results, we also hypothesize that other cues resulting in type I IFN expression, such as cytoplasmic DNA or Toll-like receptors ligands, could also depend on ILF3 to provide effective levels of these relevant antiviral products.

To date, it is still unknown how ISGs escape the host translational shutoff and engage with polysomes in the infected cells. Upon dsRNA stimulation, TOP mRNAs and other highly expressed genes disengage from polysomes. This observation raises the exciting hypothesis that disengagement of highly expressed genes from the polysomal fractions during stress or poor-translating environments, may be an essential step to allow polysomal engagement of IFNs and ISGs. Interestingly, the regulation of *IFNB1* mRNA translation was found to be specific for the NF110 isoform, which correlates with a shift in its polysomal co-sedimentation pattern upon dsRNA stimulation. The shift of NF110 to lighter ribosomal fractions during the translational shutoff leads us to hypothesize that NF110 could be supporting successful initiation of translation of *IFNB1* and specific ISGs. In agreement with previous reports, we also found PKR to be essential for efficient association of NF110 with translating ribosomes (13), which also suggests that either direct protein-protein interactions with PKR, or some basal phosphorylating activity of PKR on NF110 may be necessary to retain this factor associated with polysomes during homeostasis.

Whereas our study supports a positive role for ILF3 in promoting type I IFN and ISGs translation in cells in which cap-dependent translation is compromised, the specific mechanism by which the translation of these cytokines bypasses the translation shutoff during the antiviral response remains unknown. Although type I IFNs increased the levels of PKR, thus amplifying the translational shutoff response, they have also been shown to stimulate the translation of certain ISGs in specific cell types. Type I IFNs can activate the AKT-mTOR pathway, which in turns phosphorylates and inactivates the repressor of translation 4E-BP1 (56). In agreement, absence of 4EBP1 enhances expression of ISG15 and CXCL10 upon IFN stimulation (57). In addition, type I IFNs activate the MAPK pathways, a necessary step for efficient translation of ISG15 and IFIT2 (58). Taking all these results into account, differential translation and production of the type I IFN, IFN-β, upon ILF3 depletion could also be indirectly affecting the levels and translation of some ISGs.

In summary, we have identified ILF3 as a stimulatory factor in the establishment of an optimal type I IFN antiviral program, in which ILF3 enhances the production of IFN-β and ISGs, thus ensuring effective levels of antiviral factors in conditions where cap-dependent translation is compromised.

## Data Availability

All high-throughput sequencing data has been deposited in the GEO database, with accession number GSE130618.

## Supplementary Data

Supplementary Data are available at NAR Online.

## Funding

This work is supported by the Wellcome Trust (107665/Z/15/Z to S.M.), S. F.W. was supported by a University of Edinburgh PhD fellowship. N.B was supported by the National Research Council in Argentina (CONICET).

## Conflict of interest

The authors declare no competing interests.

## Acknowledgements

We thank our colleagues at the Institute of Immunology and Infection Research for advice and discussions. We thank Jeroen Witteveldt, Thomas Tan and Atlanta Cook for critical reading of the manuscript, and Peter Simmonds (University of Oxford) for providing Echovirus 7.

---

**Supplementary Figure 1.**
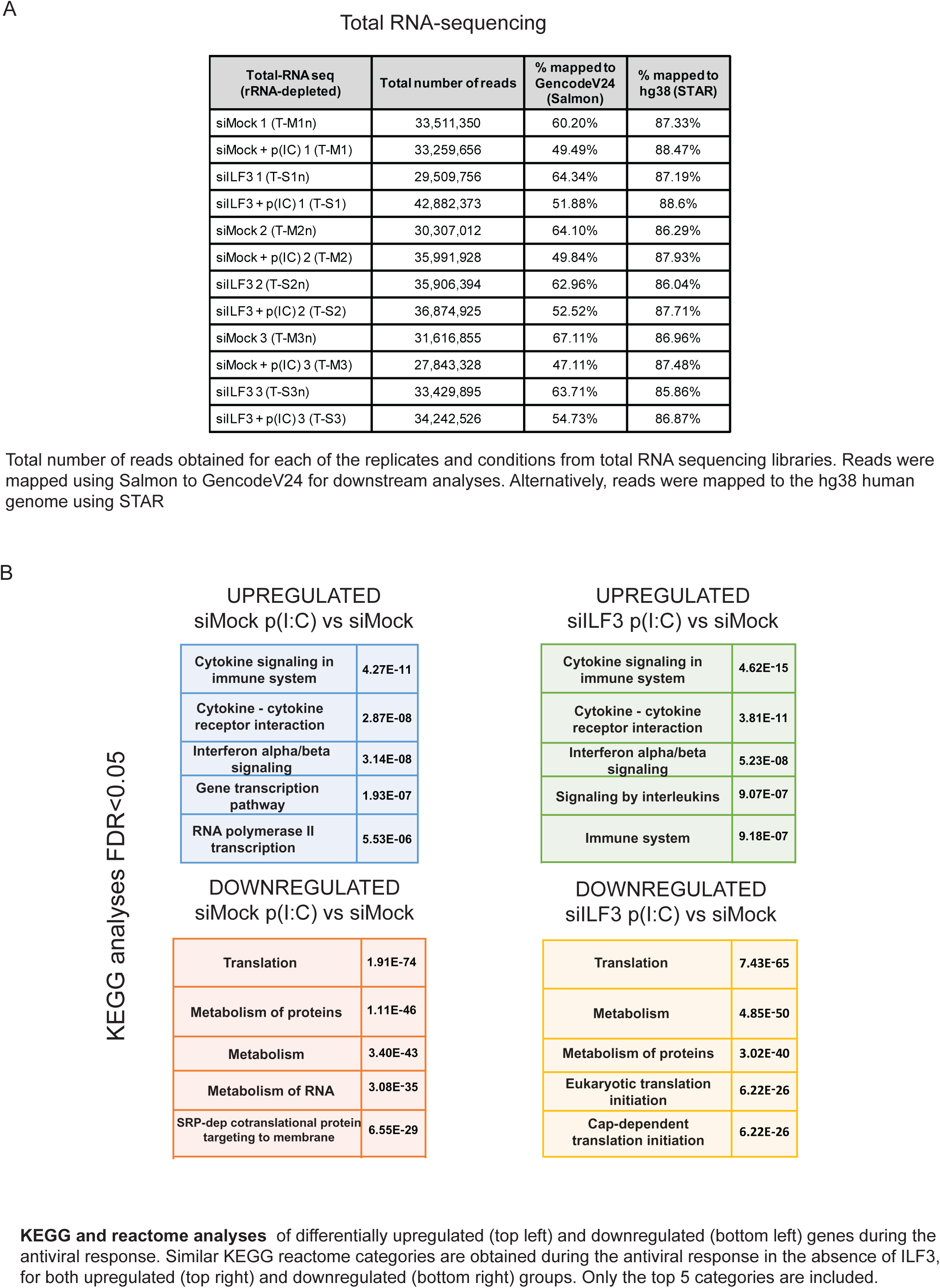

**Supplementary Figure 2.**
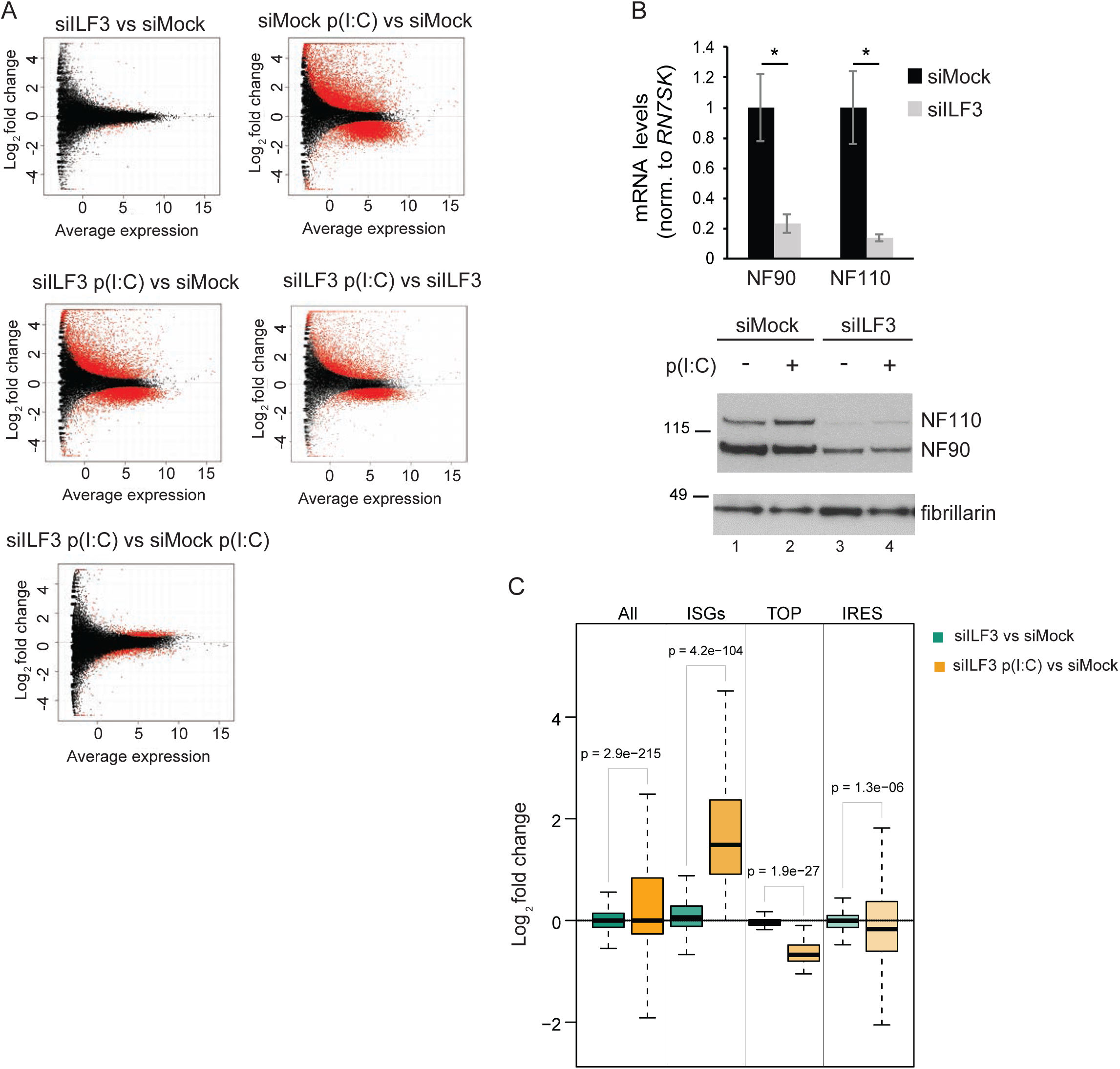
**(A)** Differentially expressed genes in red (FDR q-value <0.05) during (**1**) ILF3 depletion in homeostasis (siILF3 vs siMock), siMock represents cells transfected with non-targeting siRNAs (**2**) during the antiviral response in the presence of ILF3 (siMock p(I:C) vs siMock) (**3**) during the antiviral response in the absence of ILF3 (siILF3 p(I:C) vs siMock and siILF3 p(I:C) vs siILF3) and (**4**) ILF3-dependent genes during the antiviral response (siILF3 p(I:C) vs siMock p(I:C)). **(B) (top)** HeLa cells were depleted for 72 hours with siRNAs against ILF3 targeting both alternatively spliced isoforms produced by *ILF3* gene, NF90 and NF110. As a control, cells were transfected with non-targeting siRNAs (siMock). Isoform-specific primers were used for amplification of NF90 and NF110 isoforms to assess their relative mRNA expression levels upon ILF3 depletion. Data show the average n=3 +/− sd, (*) p-value <0.5 by Student’s T-test (**Bottom**) Representative western blot analyses of NF90 and NF110 protein levels upon depletion with siRNAs against ILF3 (lanes 3 and 4), compared to mock siRNA transfected cells (lanes 1 and 2, siMock), in the presence (lanes 2 and 4) or absence (lanes 1 and 3) of antiviral response by transfection with the dsRNA analogue, p(I:C) **(C)** Box-plot analyses of differential gene expression for: all genes, significantly induced ISGs, TOP and IRES mRNAs absence of ILF3 during homeostasis (siILF3 vs siMock, green) and in the absence of ILF3 during the antiviral response (siILF3 p(I:C) vs siMock, yellow), p-val by Mann-Whitney U test

**Supplementary Figure 3.**
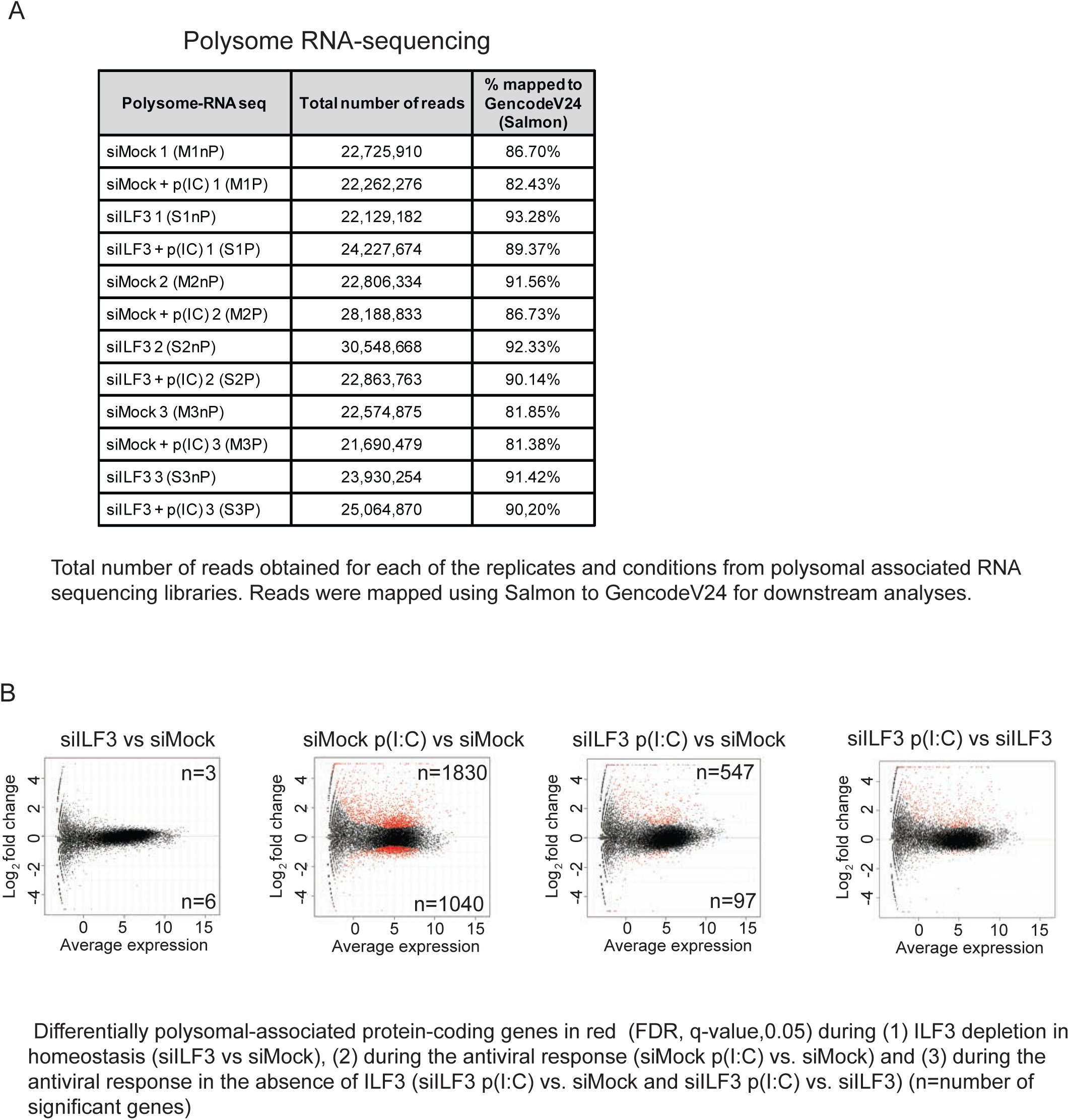

**Supplementary Figure 4.**
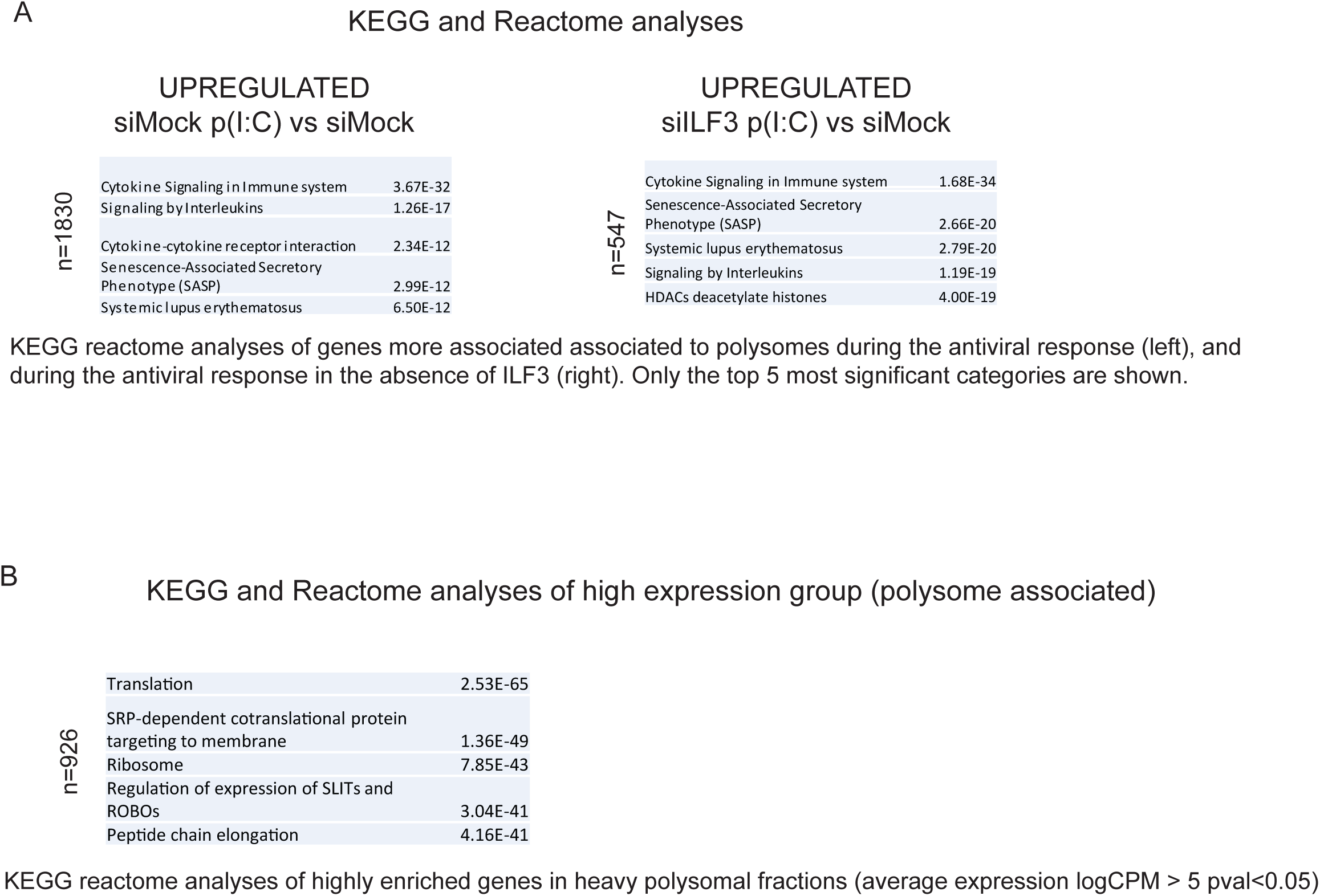

**Supplementary Figure 5.**
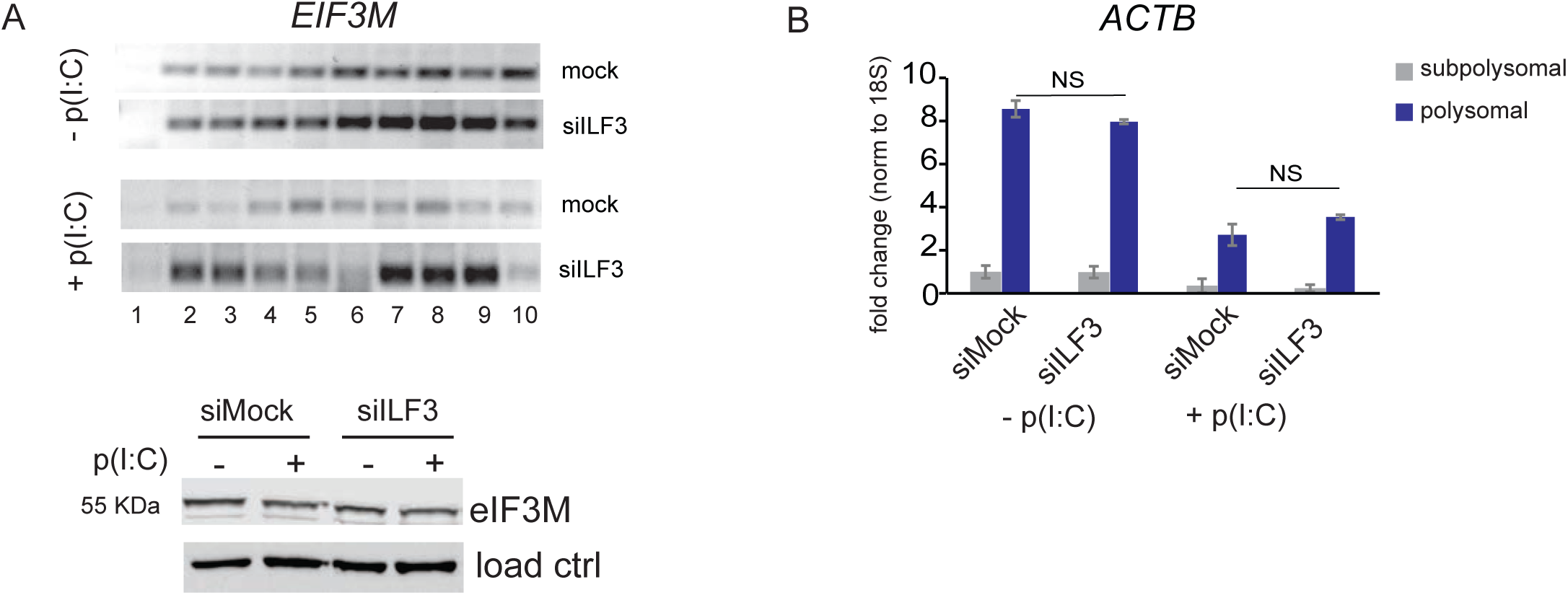
(A) (**top**) RT-PCR analyses of *EIF3M* co-sedimentation in each of the polysomal fractions collected as in Figure 3D (**bottom**) Western blot analyses of eIF3M protein levels in dsRNA-activated HeLa cells in the presence (siMock, lane 2) and absence of ILF3 (siILF3, lane 4). Fibrillarin serves as a loading control. (B) qRT-PCR analyses of *ACTB* mRNA enrichment in subpolysomal (grey) and polysomal (blue) pooled fractions in siMock or siILF3 depleted HeLa cells +/− p(I:C). Data show the average (n=2) +/− s.e.m, normalised to 18S rRNA and relative to subpolysomal levels in mock, n.s. non-significant by two-way ANOVA followed by Tukey’s multiple comparison test.

**Supplementary Figure 6.**
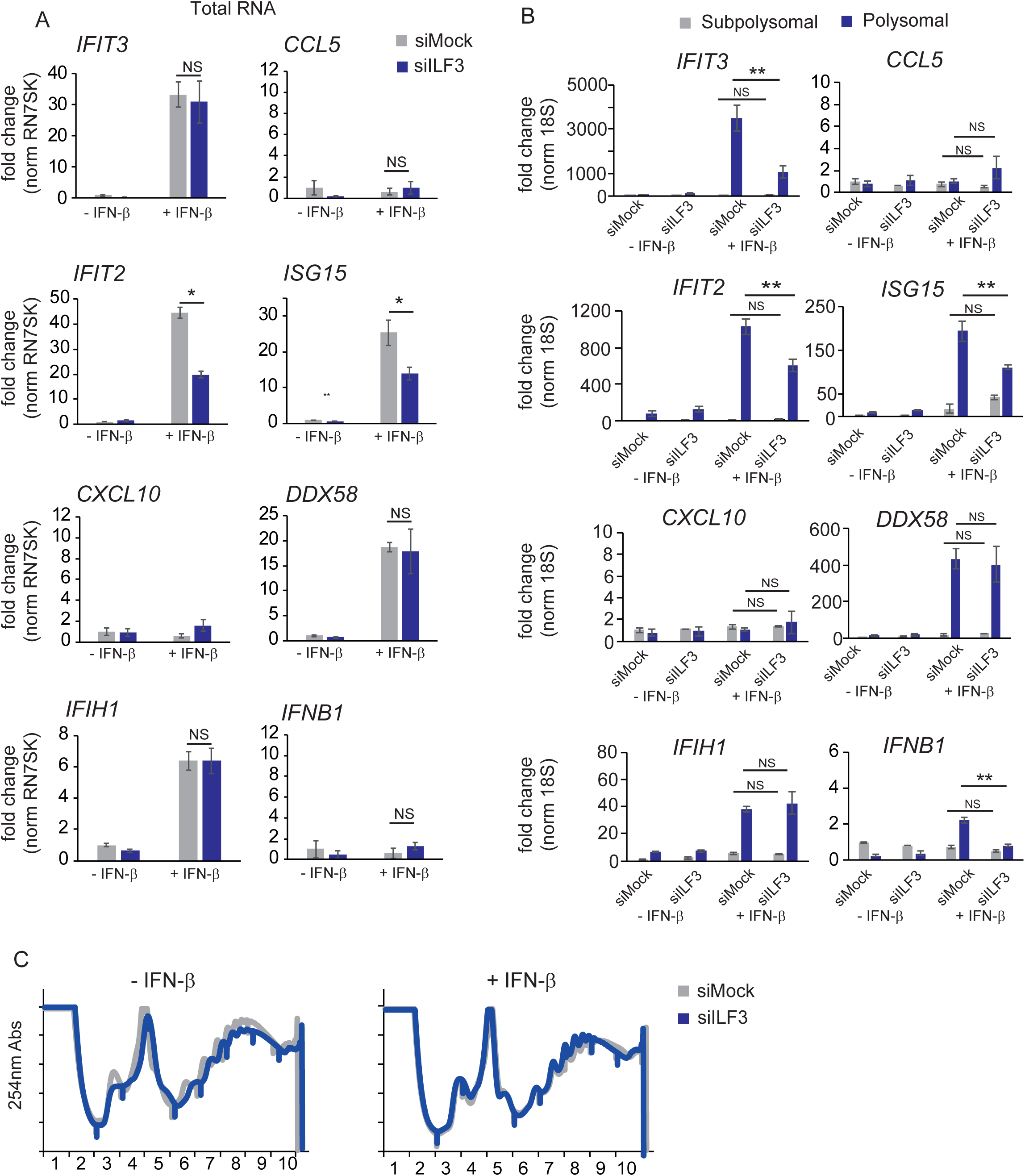
**(A)** qRT-PCR analyses of ISG and *IFNB1* expression upon 4-hour of exogenous IFN-β stimulation in HeLa cells depleted of ILF3 (siILF3, blue), a non-targeting scramble siRNA was used as a control (siMock, grey). Data show the average (n=3) +/− sem relative to (siMock -IFN-b) and normalized to RN7SK. (*) p-val<0.05 by two-way ANOVA followed by Tukey’s multiple comparison test, N.S non-significant **(B)** qRT-PCR quantification of ISGs enrichment in subpolysomal (grey) and polysomal (blue) pooled fractions with or without exogenous IFN-β stimulation, in the presence (siMock) or absence of ILF3 (siILF3). Data show the average (n=3)+/− sem normalised to 18S rRNA and relative to subpolysomal level in mock (*) p-val<0.05, (**) p-val<0.001 by two-way ANOVA followed by Tukey’s multiple comparison test, n.s. non significant **(C)** Sucrose fractionation of cytoplasmic extracts from non (-IFN-β) or stimulated with exogenous IFN-β (+IFN-β) in HeLa cells transfected with scramble siRNAs (siMock) or siRNAs targeting ILF3 (siILF3). UV absorbance, 254nm, is represented in the y-axis for each of the fractions collected after centrifugation (x-axis).

